# Data-driven signal analysis of sensory cortical processing using high-resolution fMRI across different studies

**DOI:** 10.1101/2023.08.01.551587

**Authors:** Lucas Plagwitz, Sangcheon Choi, Xin Yu, Daniel Segelcke, Esther Pogatzki-Zahn, Julian Varghese, Cornelius Faber, Bruno Pradier

**Affiliations:** Institute of Medical Informatics, University of Münster, Münster, Germany; Max Planck Institute for Biological Cybernetics, Tübingen, Germany; Athinoula A. Martinos Center for Biomedical Imaging, Department of Radiology, Harvard Medical School, Massachusetts General Hospital, Boston, USA; Department of Anesthesiology, Intensive Care and Pain Medicine, University of Münster, Münster, Germany; Department of Radiology, Translational Research Imaging Center, University of Münster, Münster, Germany

**Keywords:** fMRI, line-scanning, unsupervised machine learning, clustering analysis

## Abstract

The analysis of large data sets within and across preclinical studies has always posed a particular challenge in terms of data volume and method heterogeneity between studies. Recent developments in machine learning (ML) and artificial intelligence (AI) allow to address these challenges in complex macro- and microscopic data sets. Because of their complex data structure, functional magnetic resonance imaging (fMRI) measurements are perfectly suited to develop such ML/AI frameworks for data-driven analyses. These approaches have the potential to reveal patterns, including temporal kinetics, in blood-oxygen-level-dependent (BOLD) time series with a reduced workload. However, the typically poor signal-to-noise ratio (SNR) and low temporal resolution of fMRI time series have so far hampered such advances. Therefore, we used line scanning fMRI measurements with high SNR and high spatio-temporal resolution obtained from three independent studies and two imaging centers with heterogeneous study protocols. Unbiased time series clustering techniques were applied for the analysis of somatosensory information processing during electrical paw and optogenetic stimulation. Depending on the similarity formulation, our workflow revealed multiple patterns in BOLD time series. It produced consistent outcomes across different studies and study protocols, demonstrating the generalizability of the data-driven method for cross-study analyzes. Further, we introduce a statistical analysis that is entirely based on cluster distribution. Using this method, we can reproduce previous findings including differences in temporal BOLD characteristics between two stimulation modalities. Our data-driven approach proves high sensitivity, robustness, reproducibility, and generalizability and further quickly provides highly detailed insight into characteristics of BOLD time series. Therefore, it holds great potential for further applications in fMRI data including whole-brain task and resting-state fMRI, which can support fMRI routines. Furthermore, the analytic framework can be used for datasets that have a time-dependent data structure to integrate study results and create robust and generalizable datasets, despite different study protocols.

## 1. Introduction

Data-driven analytical techniques and decision-support systems are an active research topic in both medicine and science (Celi et al., 2019). However, for these approaches to be successful, large amounts of data from e.g., clinical trials or preclinical animal studies are needed. This challenge can only be met with difficulty by individual studies. Therefore, analytical frameworks are needed that enable the analysis of data across studies even when they were acquired independently and with different study protocols and data structures. The advantages of such approaches are valuable for microscopic (e.g., Ca^2+^ imaging, *omics data of RNA or proteins) as well as macroscopic (e.g., behavioral analyses, comparison of health data) approaches that involve complex data structures. Imaging techniques such as functional magnetic resonance imaging (fMRI) of the brain provides a showcase example of such structures with the availability of large amounts of data measured over time.

Visualization of the blood oxygen level dependent (BOLD) effect, which includes blood flow, blood volume, and blood oxygen level, allows fMRI to study cerebral processing (Buxton et al., 2004). The evaluation of these datasets can be enriched in several ways using data-driven analytical tools (Li et al., 2009). Besides the possibility of real-time analysis, there are benefits from development of new approaches through an unbiased perspective. The advantages in this regard will only be achievable if the methodology is qualitatively validated in terms of reproducibility, generalizability, robustness, and significance capability (Pineau et al., 2021). With these aspects in mind, a sub-discipline of unsupervised Machine Learning, clustering approaches such as hierarchical agglomerative clustering (HAC), are of particular interest because they build similarity relationships via a tree-like structure, the dendrogram, which can be applied to fMRI data to analyze and visualize similar activation patterns between voxels in detail. However, since similarity relations of such unsupervised analysis pipelines are susceptible to noise, fMRI data with high signal-to-noise ratio (SNR) and high temporal resolution are needed to disentangle temporal dynamics of BOLD responses.

Recent advances in small animal functional imaging enabled investigation of inter-cortical and inter-laminar processing via recording of BOLD dynamics with up to 50 ms temporal and 50 µm spatial resolution using line scanning fMRI and technical modifications thereof. This method recently demonstrated different temporal kinetics of the BOLD response in cortical laminae following sensory stimulation (Silva & Koretsky, 2002), and further, reflecting the canonical, thalamo-cortical signaling, found the BOLD response in layer IV of the cortex to precede those in laminae II/III and V (Yu et al., 2014). Furthermore, line scanning fMRI unveiled a temporal delay in cortical BOLD responses following sensory stimulation when compared to local cortical activation using optogenetic tools (Albers et al., 2018) and inter- and intra-laminar-specific patterns of functional connectivity during resting-state and electrical fore paw stimulation (Choi et al., 2022, 2023).

While advanced imaging techniques for the investigation of brain function and circuits have experienced substantial progress, the analytical tools remained similar over the recent years (Conklin et al., 2014). The typical analysis routine of fMRI time series assumes (apart from standard preprocessing) anatomical segmentation and spatial averaging of the measured BOLD signals to extract structure-specific stimulus response properties, including temporal kinetics (for e.g., onset, rise, and decay characteristics) of the BOLD response. Study results often depend on decisions of imaging experts for anatomical referencing or are derogated by spatial-averaging procedures, which severely limits their reproducibility and comparability (David et al., 2013). To abate these deficiencies, conventional analysis can be complemented and enriched by a new data-driven workflow, including different clustering approaches, that give unbiased insights into characteristics of BOLD time series.

In general, clustering analyses are used to find patterns in the datasets based on similarities (Duda & Hart, 1974). In the case of BOLD signals, these are used to detect similarities between stimulus responses within defined time periods. This technique consists of two components: (1) a similarity function and (2) a clustering algorithm, which can identify clusters and their boundaries based on the point distance (Aghabozorgi et al., 2015). The similarity function - formally known as the distance metric - defines the properties used to evaluate similarity from one observation to another. Based on these calculations, the algorithm forms groups of similar observations, e.g., similar BOLD responses.

Cluster analyses have previously been used for stimulus-induced fMRI time series. They are based on the principal observation that certain voxels can be grouped according to similarities regarding their temporal kinetics (Baumgartner et al., 1997; McIntyre & Blashfield, 1980). These similarities can be investigated focusing on frequency-based (Allegra et al., 2016) or correlation-based metrics (Craddock et al., 2012) and followed-up by numerous clustering algorithms (e.g. k-means, hierarchical, fuzzy-c means, spectral) (Cordes et al., 2002; Venkataraman et al., 2009). Problems, frequently encountered in these datasets, are (1) the large number of voxels that do not respond to the stimulation and (2) the high level of noise (Goutte et al., 1999). Importantly, these studies successfully identified stimulus-responding brain structures through unsupervised methods. However, they fell short of resolving the temporal kinetics of BOLD time series, including the rise and decay characteristics (Goutte et al., 1999).

In addition to clustering, Independent Component Analysis (ICA) is an important technique in the field of unsupervised analysis methods on fMRI data to identify patterns and structures in the data. ICA estimates independent signal sources, thereby decomposing overlapping activation patterns. Building on these sources, multiple spatial maps are created, with each map representing an independent component. These maps highlight the region exhibiting maximal activity associated with their corresponding component. In contrast, cluster analyses group similar data points together. This enables the identification of patterns in the measured data. Due to the flexibility of the similarity relations to be defined, hidden structures can be revealed depending on characteristic features such as signal strength or signal speed (Ergüner Özkoç, 2020; Korczak, 2012). While this method is strong at revealing spatial patterns of activity, it does not give insights into the temporal evolution of the BOLD time course.

The aim of this work was to assess whether the high spatial and temporal resolution of line scanning fMRI data combined with a sophisticated distance function design is sufficient 1) to detect different temporal kinetics of cortical BOLD responses and 2) to allow statistical comparison of data from different studies. By adopting a voxel-based and purely data-driven approach, we systematically explored different clustering techniques and analyzed somatosensory cortex fMRI data obtained during fore paw stimulation in rats, which were measured in different imaging centers. We established and validated an unsupervised framework that is sensitive to detect neuronal response latencies between different stimulus modalities, and which produced consistent results across different study protocols. indicating robustness, reproducibility, and generalizability of this framework.

## Material and Methods

### Animals

We used three different sets of fMRI line-scanning raw data: 1) n = 4 male Sprague Dawley rats, 2) n = 4 male Sprague Dawley rats, and 3) n = 9 female fisher rats obtained from previous studies performed at the University of Münster and the Max-Planck Institute of Biological Cybernetics in Tübingen (Albers et al., 2018; Choi et al., 2022, 2023) (Fig. 1A). All studies were performed in accordance with the German Animal Welfare Act (TierSchG) and Animal Welfare Laboratory Animal Ordinance (TierSchVersV), in full compliance with the guidelines of the EU Directive on the protection of animals used for scientific purposes (2010/63/EU). The studies were reviewed by the ethics commission (§15 TierSchG) and approved by the state authorities (*Regierungspräsidium, Tübingen, Baden-Württemberg, Germany* (study 1 and 2) and *Landesamt für Natur, Umwelt und Verbraucherschutz, Nordrhein-Westfalen, Germany* (study 3)), respectively. Rats were housed in groups of 2–3 animals under a regular light/dark schedule (12/12 h). Food and water were available ad libitum.

**Fig. 1.**
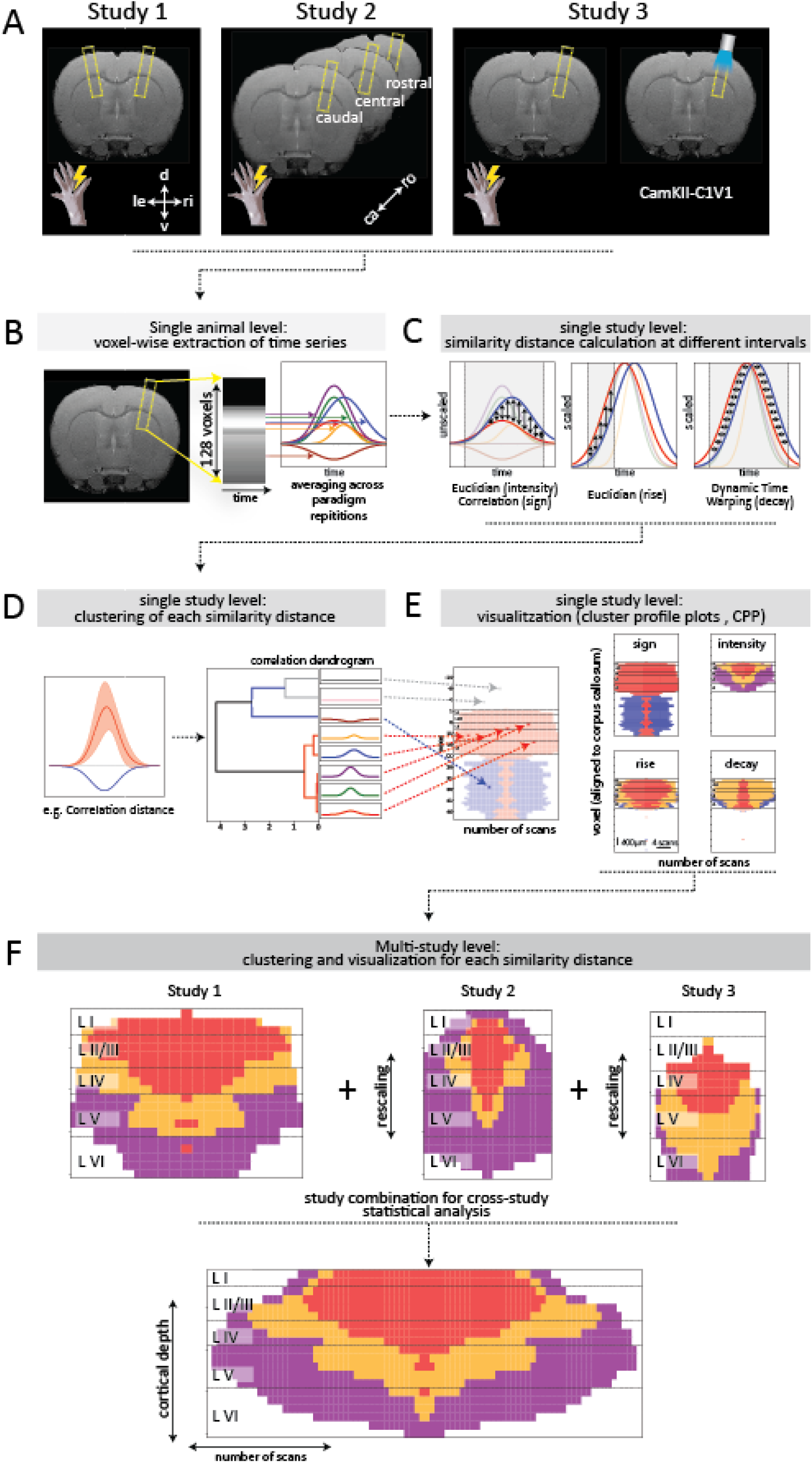
Scheme of data extraction, representation, and analysis. (A) Study designs. (B) BOLD signals are extracted and preprocessed voxel-wise from each measurement. (C) The resulting time series are compared with each other in four different settings: twice unscaled (Euclidean and Correlation) and twice scaled (Euclidean and Dynamic Time Warping (DTW)). (D) Based on the obtained distances, clusters were calculated (here only shown for correlation distance). (E) The resulting clustering distributions are illustrated with the Cluster Portability Profile (CPP), which provides the spatial location (vertical axis) and the occurrence (horizontal axis) of the individual cluster members. *Note that CPP plots are symmetric along horizontal axis.* Abbreviations: d: dorsal; v: ventral; le: left; ri: right; ca: caudal; ro : rostral; CPP : cluster probability profile.

### Line-scanning fMRI and stimulation

In table 1, recapitulates most relevant methods and parameters that were employed in the respective studies to obtain single (study 3 (Albers et al., 2018)) or multi-slice (study 1 (Choi et al., 2023) and study 2, (Choi et al., 2022)) line-scanning fMRI data (Tab. 1). Since these methods are already published, we compiled more detailed information in the supplemental material. In brief, for line-scanning fMRI, the field of view (FOV) was placed in the slice of maximal BOLD activation (Fig. 1B, left). Frequency encoding direction was set perpendicular to the cortical surface and saturation slices were used to set the width of the line to avoid aliasing artifacts. The phase-encoding gradient was turned off to acquire line profiles.

**Table 1.** Overview of different study designs and protocols. Abbreviations: TR: repetition time (ms), TE: echo time (ms), FOV: field of view, TA: acquisition time (ms), tSNR: temporal signal-to-noise-ratio

|  | Study 1 | Study 2 | Study 3 |
| --- | --- | --- | --- |
| species | Sprague-dawley rats | Sprague-dawley rats | Fisher rats |
| anesthesia | alpha-chloralose | alpha-chloralose | medetomidine |
| **Respiration** | Intubated/pancuronium | Intubated/pancuronium | spontaneously |
| **Scanner** | Magnex, Oxford | Magnex, Oxford | Bruker Biospec |
| **Field strength (T)** | 14.1 | 14.1 | 9.4 |
| **Scanner bore (cm)** | 26 | 26 | 20 |
| **# of slices** | 1 contralateral<br>1 ipsilateral | 3 contralateral | 1 contralateral |
| **TR/TE (ms)** | 100/12.5 | 100/9 | 50/18 |
| **Flip angle** | 45° | 50° | 13° |
| **FOV (cm<sup>2</sup>)** | 1.2x6.4 | 1.2x6.4 | 2.1x10 |
| **Slice thickness (mm)** | 1.2 | 1.2 | 1.2 |
| **Spatial Resolution**<br>**(μm)** | 100 | 50 | 78 |
| **Stimulation block (s)** | 4 ON 16 OFF | 4 ON 16 OFF | 5 ON 25 OFF |
| **Block repetitions** | 32 | 32 | 64 |
| **TA (min:sec)** | 10:40 | 10:40 | 32:00 |
| **Voxels per line / cortex** | 128 / 20 | 128 / 40 | 128 / 28 |
| **Cortical tSNR**<br>**over 4 seconds**<br>**(starting at stimulus)** | contralateral:<br>2.2% / 0.04% ≈ 56.9<br>ipsilateral:<br>0.005% / 0.05 % = 0.1 | Caudal:<br>2.2% / 0.03% ≈ 60.9<br>Central:<br>1.2% / 0.05% ≈ 23.7<br>Rostral:<br>0.5% / 0.1% ≈ 4.76 | Optogenetic:<br>1% / 0.03% ≈ 27.7<br>Electric:<br>0.47% / 0.03% ≈ 10.9 |

### Data preprocessing and alignment

Even though the presented approach was largely automated, the data was preprocessed with few transformation steps. First, the measurements were averaged across all repetitions, which restricted the repetitive procedure to a 20-(study 1, 2) or 30-second (study 3) signal response (Fig. 1B, right). At this point, as in previous studies (Albers et al., 2018), low-pass filtering was performed at 0.4 Hz on the averaged BOLD response to eliminate noise and to smooth the signals. In a final step, the position of the corpus callosum (CC) was determined for a later location-dependent comparison across different scans. This was done according to the profile of the average intensity over time (see Figure 1). The first local minimum in ventral direction from the global maximum was considered as the reference point of the corpus callosum. For ten measurements, no clear identification of the corpus callosum was possible, which is why these were excluded from further analyzes: Two measurements in study 1, five measurements in study 2, and three electrical stimulation scans in study 3. In study 2, the CC-detection was only possible in caudal and central layers, because of its the curvature the corpus callosum showed poor visibility in the rostral layer. In this slice, the scans were positioned based on the maximum intensity averaged over time. For all remaining 54 measurements (study 1: 20, study 2: 12, study 3: 22), the change in BOLD signal was calculated using the average of the resting period, resulting in an approximately zero-mean characteristic for the resting period.

### Clustering analysis

To partition the scans into meaningful spatial regions via a data-driven method, we followed an unsupervised machine learning approach, called cluster analysis. This technique uses the combination of a similarity metric and a clustering algorithm to detect patterns in the underlying data. Obtained clusters of similarly responding voxels are specific to the respective metric. In our approach, two types of analyzes were applied: First, similarity was analyzed in unscaled time series using the Euclidean and correlation distance (Fig. 1C, left). Second, to eliminate the influence of BOLD amplitudes and to focus on the temporal (onset and decay) characteristics of BOLD responses, the previously detected positive BOLD signals were maximum-scaled. Rise characteristics were examined with the Euclidean distance in second 1-3 post stimulus (Fig. 1C, middle), while differences in signal decay were compared with a temporally non-sensitive distance, called Dynamic Time Warping (Fig. 1C, right). This analysis pipeline was performed sequentially with line scanning fMRI data from all three studies.

### Clustering methods

All the following cluster analyzes were based on hierarchical agglomerative clustering (HAC), which operated on predefined subsections of the respective time series (Fig. 1D). We applied this type of clustering procedure since it is less sensitive to largely differing point densities of clusters, and it can represent nested clusters (Sander et al., 2003). Ward’s minimum variance method was used for the optimization process within the cluster formation (Ward, 1963). For the application of HAC, Python was used with the Scikit-Learn package (Pedregosa et al., 2011). In addition to the different time intervals, three different metrics were considered as a basis for similarity relationships between two time series x, y ∈ ℝ*^n^*^≤600^ with length n:

1. Euclidean distance

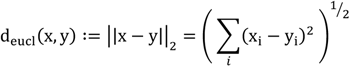

1. Correlation distance

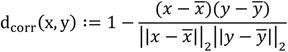

1. Dynamic Time Warping (DTW)

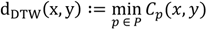

where *C_p_* denotes the costs for a warping path *p* from *x* to y.

These metrics were used at different time intervals to highlight different properties of a time series.

### Characterization of unscaled BOLD-signal change

For a first overview of the measurements, the dataset was restricted to the range of the approximate stimulus response (time of stimulus onset until twice the length of the stimulus; study 1, 2: 8s, study 3: 10s). Thus, the cluster membership was determined for each of the series and their corresponding voxels. First, Euclidian distance (d_eucl_(x, y)) and correlation distance (d_corr_(x, y)) were considered for a basic comparison of similarities. Subsequently, the computed clusters were characterized by their point-wise mean and standard deviation. In addition, we developed a new type of visualization called the Cluster Portability Profile (CPP) plot to highlight the spatial distribution of patterns identified in 1D fMRI measurements. For this purpose, the described line-centering based on the CC was performed (see above). Based on this centering, the expected size of the cortex was calculated and divided into five parts according to cortical layers: LI (CC-[210 - 190 µm]), LII/III (CC-[190 - 150 µm]),), LIV (CC-[150 - 120 µm]),), LV (CC-[120 - 70 µm]),) and LVI (CC-[70 - 10 µm]). Boundaries of these layers were marked with dashed lines in the CPP plot. Statistical tests were performed based on this division. To validate the alignment of 1D lines for the CPP plot, we compared the automated approach to manually drawn cortical boundaries in 2D BOLD fMRI data. The stability of the method was assessed using animal-based 5-fold cross-validation (McIntyre & Blashfield, 1980). This involved repeated clustering of a subset of available animals. For this test, we used data from study 3 since it included the largest number of animals. Performance was then quantified using three quality measures: V-measure, sensitivity, and specificity (Rosenberg & Hirschberg, 2007). The clusters that showed a sizable positive BOLD response were used to define cortical pixels. These were compared to the manually determined cortical boundaries.

### Time series clustering based on the rise and decay characteristics of BOLD activation

The next step focused on the kinetics of the stimulus response, including the rise and decay characteristics of the BOLD signal. For this purpose, we extracted time series from the cluster with the strongest response as detected by the correlation-based cluster analysis. Next, to avoid noise amplification, we added a criterion of a 0.5% minimum signal change. All signals, which fulfilled these two characteristics (highest cluster and ≥ 0.5% BOLD change), were scaled by their maximum. This subset was then analyzed in two different ways: (1) by Euclidean clustering in the interval 1s-3s post stimulus onset for the characterization of the BOLD signal rising period and (2) by DTW over the entire previously considered intervals for the decay kinetics. In the latter case, the temporal onset was neglected by observing the non-time sensitive DTW. The analysis of the resulting findings followed the same structure as described above, with characterization based on the time-dependent mean and standard deviation and a CPP plot by type of stimulus (electrical vs. optogenetic) and study. All three studies were analyzed with this workflow and produced consistent results, indicating generalizability of this workflow. Comparably, this rise and decay analysis, as well as the previous assessment of the non-normalized time series, was performed with the dataset of the second study. The consistency of the study protocols allowed the use of appropriate time intervals.

### Layer-dependent response analysis across multiple studies

In a final step, we examined the signal processing of electrical paw stimulation in terms of intensity, rise, and decay across all three studies. For this purpose, the cluster-based parameters within each study were calculated, using only those of the caudal slice in study 2 and omitting optogenetic measurements in study 3. As an additional adjustment for the intensity analysis, the time series of a measurement were scaled by the respective measurement-specific maximum signal response. To allow cross-study analysis, spatial resolution of the 1D lines had to be corrected to the same length. To account for the different spatial resolutions (see Table 1), and hence to ensure comparability of cortical dimensions across all studies, we performed a voxel transformation. For this purpose, all voxels from study 2 and 3 were aligned to the size of study 1. In study 2, this involved a process of compressing 40 voxels into a streamlined group of 20 though a pairwise averaging technique. For study 3, we reconfigured the original 28 cortical voxels to match the 20-voxel structure of study 1. To this end, we multiplied each voxel 5 times, resulting in a cortical size of 140 voxels. Next, we average groups of 7 voxels each, thereby effectively reducing the voxel numbers to our target size of 20 voxels per cortex. This guaranteed a comparable size for the CPP plots. In the procedure, the combination of ordinal-scaled clusters with each other was possible, even if the study protocols did not fully match. This resulted in CPP plots with a high number of measurements so that a layer-based significance analysis could be performed.

### Statistical Analysis

We tested different stimuli (electric vs. optogenetic) and regions (cortical layers) for significant differences with respect to cluster elements. For this purpose, the clusters were first sorted in ordinal order, so that there was an ascending sequence of cluster groups. Depending on the extraction property, these time series were ordered by intensity, rise, or decay of activation from low to high or slow to fast, respectively. Subsequently, the median value of this order was determined for each measurement. Thus, distributions were obtained across the measurements, which were compared against other conditions. Differences between electrical and optogenetic are determined using the Mann-Whitney U rank test, while layer-based comparisons are calculated using Wilcoxon signed-rank test.

## Results

### Data-driven detection of cortical boundaries and clustering of BOLD signal properties

First, we analyzed similarities between the contralateral time series of study 1 following electrical fore paw stimulation (Fig. 1A, left). Using the correlation metric in an interval covering the whole BOLD response, averaged across 30 repetitions, with a predefined number of three clusters, we found signals with a positive (red) or negative (blue) characteristic, contrasting a third neutral cluster (white, Fig. 2A, left). These clusters were spatially segregated into voxels that were dorsal (red) or ventral (blue) to the corpus callosum (CC) as shown in the CPP plot, thereby providing the possibility to discriminate cortical from sub-cortical (i.e., striatal) clusters (Fig. 2A, right). On the ipsilateral hemisphere, the BOLD percent change in time series were significantly smaller (< 0.1% change) and could not be assigned to cortex- or striatum-specific clusters, indicating the SNR as critical variable for this analysis (Fig. 2B). Given that averaging improves SNR, we sought to explore how this would impact the performance of clustering on contralateral time series. To do so, we conducted the clustering process using a reduced number of repetitions for averaging. Despite the lower number of repetitions (as few as 4-16), we found that this approach was sufficient to yield stable clustering results in both the cortex and the striatum (Fig. 2C). This finding suggests that stable results can be achieved with fewer repetitions and lower SNR. Next, using the Euclidean distance we detected four clusters on the contralateral hemisphere: three of them in the cortex, which were characterized by amplitude, full width at half maximum (FWHM) and undershoot (purple < yellow < red), and contrasted with a neutral (white) fourth cluster (Fig. 2D, left). Because these clusters were characterized by different properties of the BOLD signal, we described them with the general term intensity level. We observed a gradient in the cluster distribution along the dorso-ventral axis in the S1FL cortex (Fig. 2D right): voxels of the high-intensity cluster (red, dorsal-most) were followed by those with medium-(orange, dorsal-central), and low-intensity clusters (purple, ventral-most). On the ipsilateral side, we observed a similar effect as before with no clear region assignment possible, which is why we excluded these data from further analysis (Fig. 2E).

**Fig. 2.**
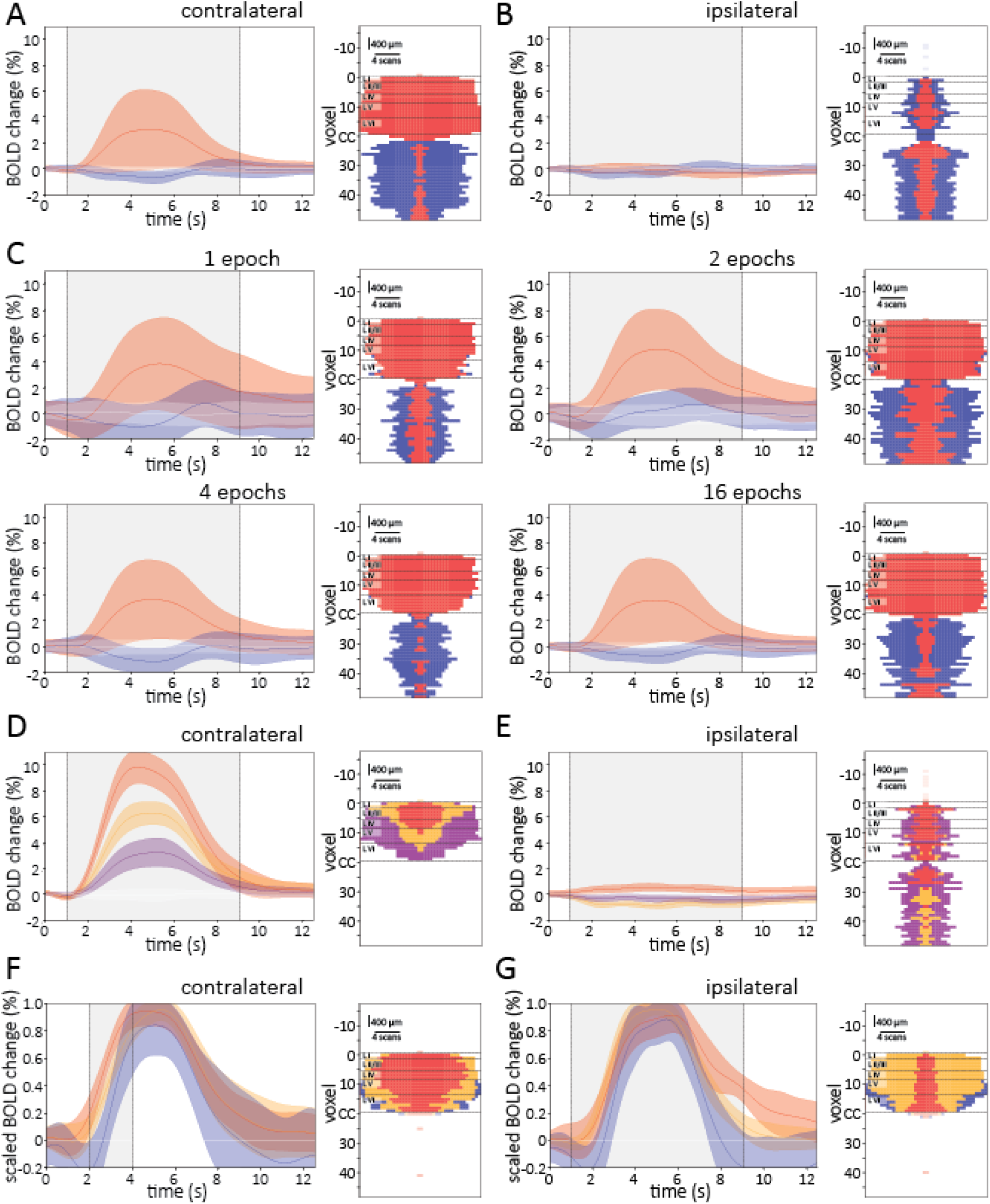
Hierarchical clustering combined with different metrics reveal spatial-dependent properties of BOLD signals at a glance. To visualize the properties and spatial distribution of different clusters, averaged time series and the CPP plot of the corresponding clustering metric are shown on the left and right of each subfigure, respectively. (A) Clustering based on correlation distance for contralateral and (B) ipsilateral line scanning data. (C) Cluster results of the correlation analysis using single epochs, or averages of 2, 4 or 16 repetitions. (D) Euclidean-based clustering of contralateral and (E) ipsilateral BOLD signals. (F) Clustering distribution and properties of the rise level-based examination based on the interval 1s - 3s post stimulus onset for data from study 1. (G) Clustering result for DTW-based decay characterization. Time series are represented as mean +/− SD for each cluster, gray-shaded area represent time interval used for cluster calculation. To visualize the spatial representation of cluster distribution in the cortex, the CPP plot of the corresponding clustering is shown. Note that CPP plots are symmetric along horizontal axis. *Abbreviations: CPP: Cluster Probability Profile, DTW: Dynamic Time Warping, SD: Standard Deviation*

To acquire information on rise kinetics of BOLD responses with a positive sign in the cortex (see red cluster Fig. 2A), we performed Euclidian-based clustering on scaled time series during the interval 1–3 s following the stimulation (Fig. 2F, left). Properties that characterized the clustered time series included rise time (10–90%), τ (time to half-maximum), onset, and undershoot prior to rising phase; hence we describe these clusters with the general term rise level. Red clusters collected high rise levels, orange clusters represented medium and blue delayed rise levels. Like before, we detected a gradient in the dorso-ventral axis of the cortex with the fastest clusters (red) in the dorsal-most area of the cortex (Fig. 2F, right). Next, we investigated the decay characteristic of BOLD signals using the DTW distance. This metric was used intentionally since it neglects the previously considered differences in the rise characteristic. We obtained three clusters that operated independently of the signal start using the same cortical cluster as for the analysis of rise level. These clusters segregated during the second half (5-10s post stimulus onset) of the considered time interval (gray shaded area) according to their decay characteristics (Fig. 2G).

### Validation of analysis framework on independent study

To validate our workflow, we analyzed an independent study (study 2, Fig. 1A, middle), which was performed with different acquisition parameters (see Table 1). While the temporal resolution was the same, it had a higher spatial resolution. Additionally, 3 lines were acquired simultaneously on the contralateral side. These lines extended in rostral direction with the caudal-most line starting in the same anatomical position as in the previous study (Choi et al., 2022). In the caudal line we observed similar cluster distributions as described before regarding the sign of BOLD signals (correlation-based, Fig. 3A) and intensity levels (Euclidian-based, Fig. 3B). We also obtained similar clustering results when investigating rise (Fig. 3C) and decay (Fig. 3D) characteristics, thereby validating our previous results. Further, this cluster distribution proved to be robustly present also in the central of the three lines: we found clusters with a positive sign in the cortex, and clusters with high intensity and rise levels were segregated along a dorso-ventral gradient, while those characterized with slow decay kinetic were segregated along a ventro-dorsal gradient within the cortex. In rostral lines, we recovered fewer signals and observed a further cluster (turquoise) with a negative sign with the correlation-based clustering. This cluster was found in the cortex rostral to S1FP and showed delayed onset characteristics compared to the negative BOLD responses typically found in the striatum (blue) (Amirmohseni et al., 2016).

**Figure 3).**
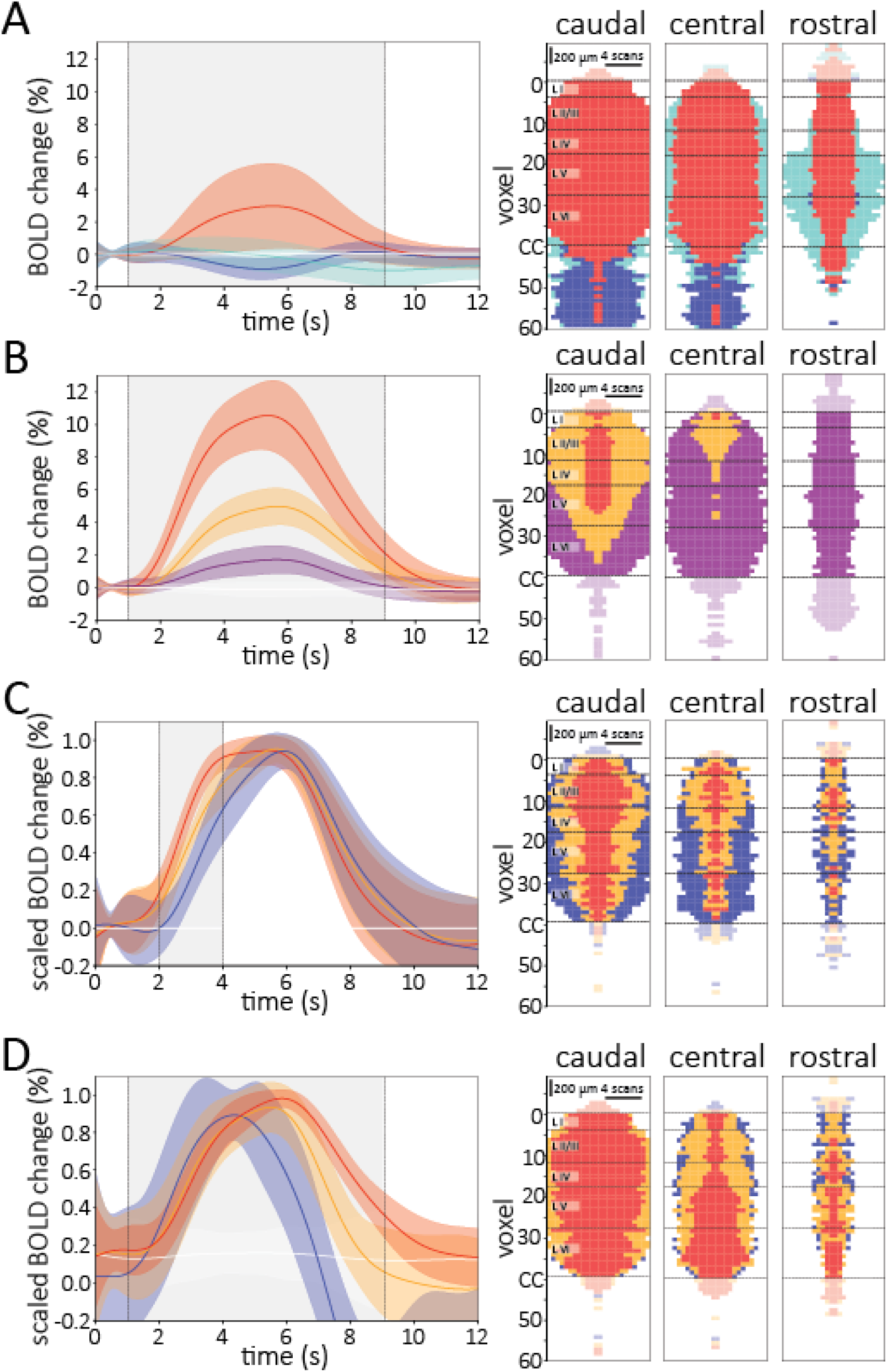
Validation of hierarchical clustering-based workflow on independent study. (A) Correlation-based clustering calculated on the interval between seconds 1–9 following stimulus onset. (B) Euclidean-based cluster distribution on the same interval in different slices. (C) Rise characteristic revealed by the Euclidean distance in the time interval 1-3s post stimulus. (D) DTW-based clustering to calculate the BOLD decay level following seconds 1 - 9. Time series are represented as mean +/− SD for each cluster, gray-shaded area represent time interval used for cluster calculation. To visualize the spatial representation of cluster distribution in the cortex, the CPP plot of the corresponding clustering is shown. Note that CPP plots are symmetric along horizontal axis. Abbreviations: CPP: Cluster Probability Profile, DTW: Dynamic Time Warping, SD: Standard Deviation

### Within-study statistical comparison of cluster distributions

Having established a workflow for a qualitative assessment of BOLD characteristics of line-scanning fMRI data, we explored the possibility to perform statistical analyzes based on cluster distributions. To this end we compared two stimulus modalities (electrical fore paw stimulation and optogenetic cortical stimulation, study 3, Tab. 1 and Fig. 1A right) between the cortical layers. Both metrics, Euclidean and correlation, operated on unscaled BOLD signals, showed similar cluster distribution for sign and intensity level of the BOLD response as described above (Euclidean: Fig. 4A and correlation: Fig. 4B). Statistical analysis revealed that intensity levels were significantly higher in layer IV, V upon optogenetic compared to the electrical stimulation (Mann-Whitney U rank test, p < 0.05, Fig. 4B). Additionally, time series following optogenetic stimulation showed clusters with faster rise levels (red) compared to electrical stimulation (Fig. 4C). This effect was significant in layer V (Mann-Whitney U rank test, p < 0.05, Fig. 4C) but not in layers II-IV (p < 0.1). Finally, when analyzing the decay characteristic, we found the red cluster with the slowest decay characteristic almost exclusively in data following optogenetic stimulation. The cluster distribution showed a significant difference in layers V and VI between electrical and optogenetic stimulation (Mann-Whitney U rank test, p < 0.05, Fig. 4D). Together, our workflow performs a robust analysis and visualization of different BOLD characteristics producing consistent results across different study protocols and further allows quantification of cluster distributions in a data-driven way.

**Figure 4).**
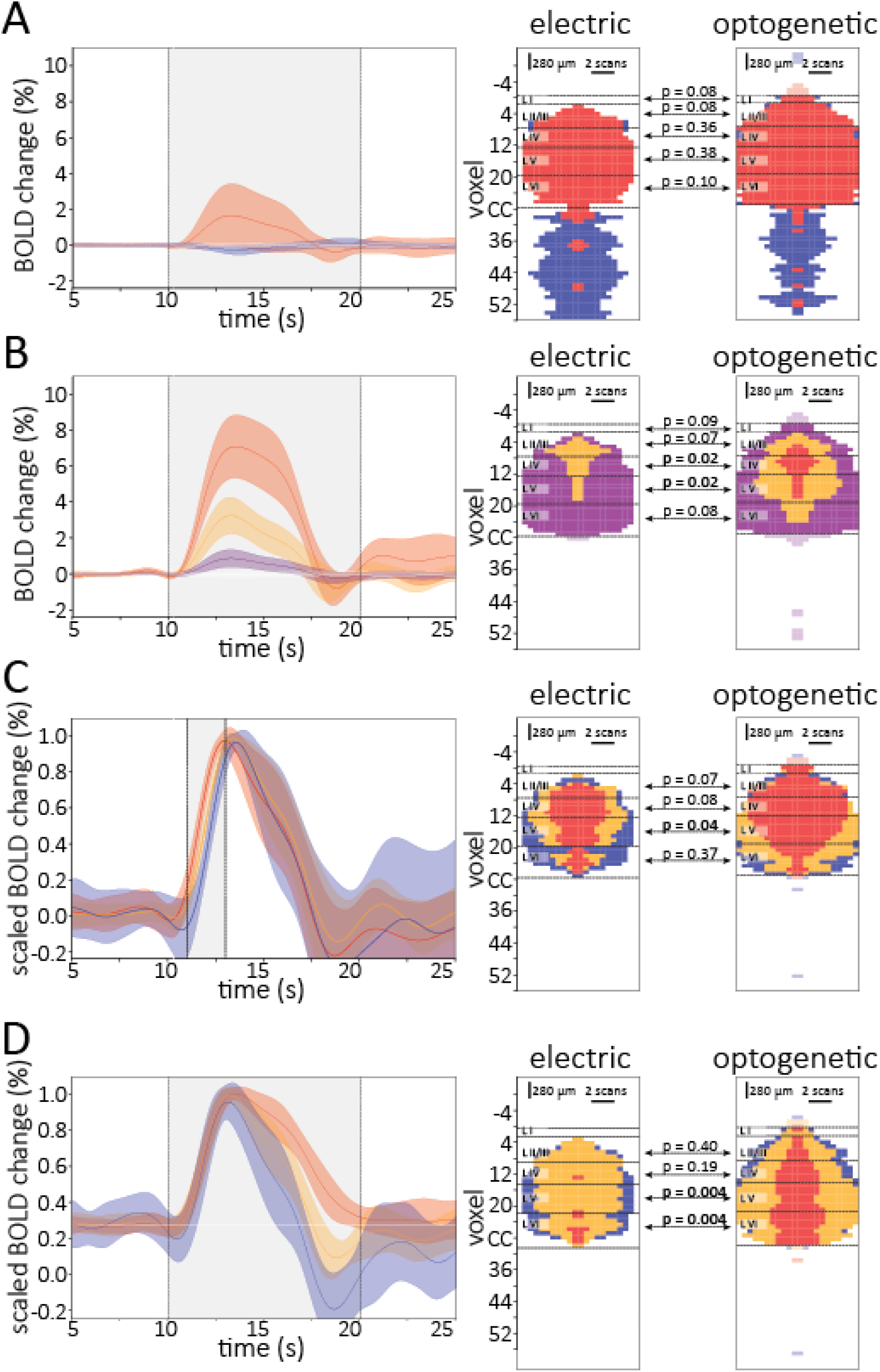
Statistical comparison of cluster distributions to analyze differences in signal characteristics between different stimulation modalities. (A) Clustering-based on correlation distance combined with statistical comparison between both modalities. (B) Result of Euclidean-based clustering of BOLD signals following electrical (middle) and optogenetic stimulation (right). BOLD signals have significantly higher intensity clusters in layers IV and V following optogenetic stimulation (Mann-Whitney-U test). (C) Euclidean-based clustering during 1s - 3s post stimulus onset shows more voxels with a higher rise level following optogenetic stimulation in layer V. (D) DTW-based clustering for differences in signal decay show clusters with slower BOLD decay kinetic following optogenetic stimulation in layers V and VI. Time series are represented as mean +/− SD for each cluster, gray-shaded area represent time interval used for cluster calculation. To visualize the spatial representation of cluster distribution in the cortex, the CPP plot of the corresponding clustering is shown. Note that CPP plots are symmetric along horizontal axis. Abbreviations: CPP: Cluster Probability Profile, DTW: Dynamic Time Warping, SD: Standard Deviation

### Stability testing of automated data alignment

In general, clustering algorithms always detect clusters, even in random data(McShane et al., 2002). Therefore, in addition to the validation of our workflow over different datasets, we were also interested to investigate the stability of the method itself. To do so, we subsampled animal-wise our input dataset and examined the behavior of unsupervised data analysis on the output. Using stability testing (5-fold cross-validation), we explicitly compared the automated alignment of 1D lines and the subsequent definition of cortical boundaries that is used to generate CPP plots with manually-annotated boundaries via mapping 1D line onto 2D anatomical images (Tab. 2). The correlation-based clustering showed high overlap to manually defined regions as revealed by overall highest values for sensitivity (0.842 ± 0.032), specificity (0.977 ± 0.008), and V-measure (0.626 ± 0.035), which is defined as the harmonic mean between homogeneity (the cortex contains only points which are member points) and completeness (all member points are elements of the cortex), and which commonly used to evaluate the correctness of clusters. When we combined all three positive clusters identified from the Euclidian metric (red, orange, and purple), a smaller portion of the cortex was tagged, causing a decrease in sensitivity to a value of 0.668. On the other hand, we saw a small increase in specificity, up to 0.988. Considering the standard deviation of the cortical area for the two metrics, correlation-based clustering results demonstrate notably increased stability. When comparing both metrics, changing input data the Euclidean-based approach was more influenced by the input data, while the correlation-based clustering stayed almost the same, even when with small input changes due to false-positive alignments outside the cortex. Together, these data show that the data-driven approach to detect cortical boundaries with either metric correlates well with those that were drawn manually. To conduct a more extensive analysis of the cortex, we opted to utilize the annotations garnered from the correlation-based clustering. This choice was informed by our observation that this particular methodology displayed the most robust alignment with the cortical structures, as evidenced by our prior examination of relevant performance metrics.

**Table 2.**
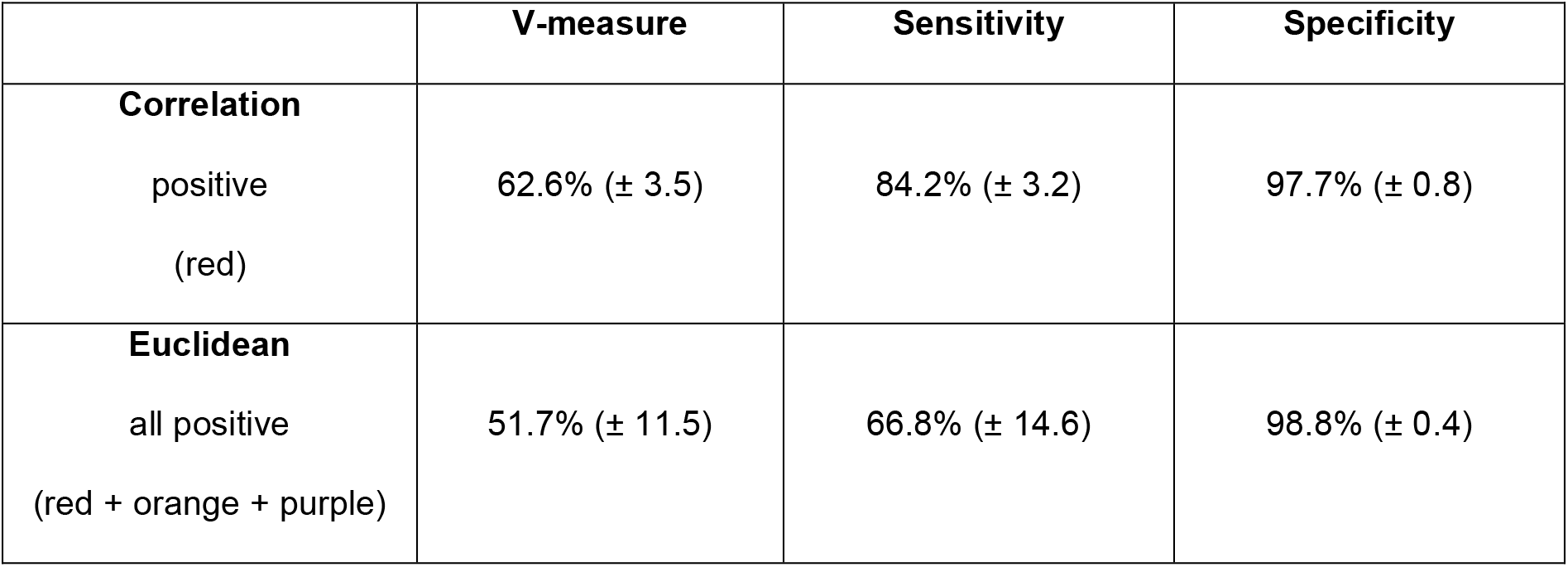
Cross-validated performance metrics for analyzing stability of automated alignment by comparing clustering results to an expert-drawn cortex annotation. V-Measure: harmonic mean between homogeneity (the cortex contains only points which are member points) and completeness (all member points are elements of the cortex), sensitivity (true positives vs. condition positives), specificity (true negatives vs. condition negatives)

### Cross-study comparison of BOLD dynamics in between cortical layers

Finally, we combined 43 data sets with electric paw stimulation across three studies and investigated the processing of sensory information between neighboring cortical layers (LI, LII/III, LIV, LV, and LVI). To assess the comparability of our results with the previously published analyses of the same data, we extracted single parameter rise characteristics of the BOLD signal, including time to 10% or 50% of maximal response amplitude (T_10_ and T_50_, respectively), in LII/III-LIV and LV-LVI (Suppl. Fig. 1). In all studies, both parameters were in similar ranges as described before (Albers et al., 2018; Choi et al., 2022, 2023) and show significantly faster BOLD signals in LII-IV compared to LV-VI. Furthermore, we found no cross-study differences in these characteristics for LII-IV and V-VI, respectively. Next, we found significantly higher intensity and rise levels when comparing LIV to LV and LV to LVI, respectively (Fig. 5A, 5B, Wilcoxon signed-rank test, p < .05). Differences between LI and LII/III were only significant at the intensity level. General systematic layer-based differences were not detectable via the decay characteristics (data not shown, Wilcoxon signed-rank test, p > 0.05) and were hence excluded from the next analysis step. Next, we investigated the correlation between rise and intensity level clusters of the respective layers. For this purpose, we considered means (stars) and covariances in the 2D plane between intensity and rise level for each layer and displayed the double standard deviation as ellipsoids (Fig. 5C). A linear dependence between rise and intensity behavior of the BOLD response is visible and a separation of LVI from LV, while LII/III and LIV showed large overlap. Furthermore, we found large agreement between clustering results from the different studies indicating successful harmonization of different study protocols and data structures through inner-study scaling (Fig. 5D).

**Figure 5).**
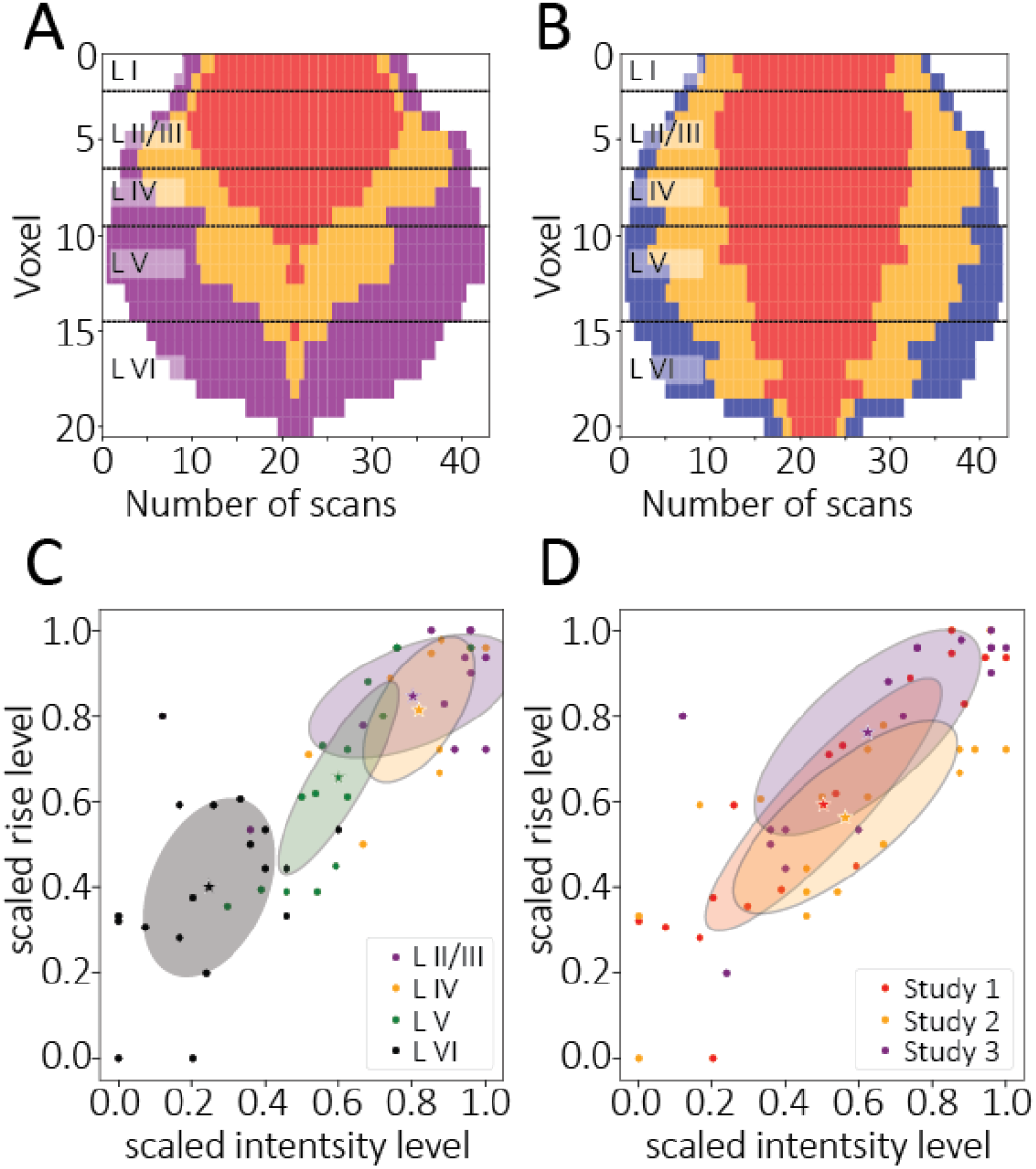
Layer-wise comparison of intensity and rise level across all three studies. (A) CPP plot of intensity levels based on non-scaled BOLD signals across all three studies. (B) Rise characteristic clustering based on time interval 1s-3s post stimulus. (C) Covariance analysis of 2D plane dependent on intensity and rise level grouped by layers. A point represents a voxel averaged over all measurements in a study. This led to 20 voxels x 3 studies = 60 points. The 4 colored stars highlight the grouped mean. (D) Covariance analysis of 2D plane dependent on intensity and rise level grouped by studies. (A) and (B) also indicate the statistical tests for dissimilarity between the neighboring layers (Wilcoxon signed-rank test). Star indicates cluster mean, ellipsoids indicate double SD. Note that CPP plots are symmetric along horizontal axis. Abbreviations: CPP: Cluster Probability Profile, SD: standard deviation.

## Discussion

The method of line scanning fMRI has already demonstrated the ability to resolve differences of a few tens of milliseconds in the BOLD onset due to its high data resolution (Yu et al., 2014). Building on these advantages of BOLD signals with high spatial and temporal resolution, we established a novel clustering analysis that can spatially separate and visualize different temporal characteristics of extracted BOLD time series and allows for mapping those onto cortical boundaries. Based on this clustering we further introduced a statistical framework that relies on multi-feature analysis of BOLD responses rather than extraction of single BOLD parameters. Using datasets from three published line scanning studies from two different imaging centers, we analyzed the influence of multiple clustering approaches to derive differently responding regions in the somatosensory cortex with high spatiotemporal resolution. Within our analysis framework, we introduced a novel approach for cross-study statistical analysis of fMRI data, based on distinct cluster characteristics, providing new insights into cortical stimulus processing in a visually compelling way. This pipeline allows existing research to be rescaled, compared, and potentially combined across studies to identify new aspects in a data-driven manner.

### Different clustering analyzes robustly detect inter- and intra-laminae-specific BOLD properties

Using the correlation-based HAC centered on the entire stimulus duration we were able to discriminate cortical boundaries from the striatum and from voxels outside the brain with high accuracy, a pivotal first step in our data-driven analysis pipeline. Then, applying the Euclidean metric on the same interval, we showed in each of the three studies a consistent separation of clusters depending on their BOLD signal intensity level. The time series with the highest intensities occurred in the dorsal-most regions of the primary somatosensory cortex, which confirms previous reports (Jung et al., 2021; Tian et al., 2010; Yu et al., 2014) and is characteristic for gradient echo fMRI data, which are more sensitive to blood oxygenation changes in larger blood vessels that are found on the cortical surface (Báez-Yánez et al., 2017; Uludağ & Blinder, 2018). In study 3, we directly compared cortical optogenetic activation of CamKII excitatory neurons with electrical fore paw stimulation by analysis of the rise and decay levels of positive time series using DTW and correlation metrics. We found faster rise characteristics in LIV and slower decay properties in LIV and LV following optogenetic compared to electrical stimulation. This is consistent with our recently published analysis (Albers et al., 2018). However, in the previous analysis this effect was detected using a hemodynamic response-fitting (HRF) procedure combined with extraction of the onset (T0) of the BOLD response. Yet, these fitting procedures are partially susceptible to bias (e.g., noise in time series) and do not account for events such as the initial dip that is occasionally reported by us and others (Uludağ & Blinder, 2018). In contrast, our rise-level clustering approach does not require fitting, but instead accounts for the entire rising phase of the BOLD response, thereby including slope and rise times of the BOLD signal ranging from T_0_ to T_max_ and can therefore be considered more holistically.

Investigating intra-laminar BOLD effects (study 2), we detected that the probability of high intensity BOLD signals clearly increased from rostral to caudal direction, which has not been reported in our previous study (Choi et al., 2022). This clearly puts the center of activation to sensory fore paw stimulation in most posterior slices, which is consistent with the topography of the area representing the foot pad of the primary sensory cortex of the fore limb (S1FL) (Chapin & Lin, 1984; Seelke et al., 2012). Positive BOLD changes in the central slice have lower intensity levels since approx. half of the slice volume is represented by digit-responsive areas of the S1FL and hence should not be activated by the stimulation of the foot pad (Chapin & Lin, 1984). Moving rostrally, the positive, yet low-intensity, BOLD responses likely reflects activation of the secondary motor cortex of the fore paw, which neighbors the S1FL in anterior direction (Ebbesen et al., 2018; Seelke et al., 2012), thereby showing intra-cortical stimulus processing and signal integration. Furthermore, consistent with our previous data analyzes, we also detected two types of negative BOLD responses in different regions: a fast-peaking response that was mostly located ventral to the corpus callosum in the striatum and which has been associated with neuronal activation (Amirmohseni et al., 2016; Shih et al., 2012). Further, a delayed-peaking response that was found in approx. half of the animals in cortical layers LV and LVI in the rostral-most slice (discussed in (Choi et al., 2022). Together, we have characterized the BOLD response depending on metric and time interval in more detail through their global properties such as intensity, rise, and decay, and thereby confirm results from previous studies with an independent and completely data-driven approach.

### Limits in detection of laminar-specific rise characteristics of BOLD signal

Recent reports segregated onset dynamics of BOLD signals following paw stimulation in the thalamocortical input layer (LIV, earliest onset), from those with delayed onsets in the adjacent layers LII/III or LV (Silva & Koretsky, 2002; Yu et al., 2014), which reflects canonical signal processing in the cortex. Therefore, we investigated whether we could resolve these differences as well and combined datasets with the same stimulation modality from all studies to obtain a global study-independent signal characterization. To allow comparison of our data with the literature we extracted rise time characteristics, including T_10_ and T_50_ and observed similar rise characteristics as described in previous reports (Albers et al., 2018; Silva & Koretsky, 2002; Yu et al., 2014). There were no differences between the studies in T_10_ and T_50_, but across all studies, we observed significantly faster BOLD time series in LII/III+IV compared to LV-VI. However, we and others (Jung et al., 2021) did not detect a significantly earlier onset in LIV compared to LII/III. While this seems contradictory to earlier findings from other studies (Yu et al., 2014), it is important to mention that the conclusions were reached with different analyzes: in this study we focused on an interval enclosing the entire rise time, whereas in the original description of the line scanning method the onset (T_0_) of the BOLD signal was estimated (Yu et al., 2014). Furthermore, the majority of our data (x out of 43, obtained from studies 1 and 2) were acquired with a temporal resolution of T_R_=100ms, which is well below the T_R_≤50ms that was used to detect inter-laminar-specific processing (SILVA, 2002, Yu 2014). Therefore, if smaller latencies in BOLD signal should be investigated faster acquisition schemes with high temporal resolution (≤ 50 ms) are necessary to detect early thalamic synaptic input and neuronal activation in layer IV, which is best combined with immobilization and mechanical ventilation of animals(Yu et al., 2014). Furthermore, the technical development of even faster fMRI sequences will help to resolve and study these differences.

### Potential of clustering analyses for fMRI time series

Previous studies have demonstrated the added value of cluster-based analyses in fMRI time series. Here, we go one step beyond distinct simultaneous activation analysis and add voxel-based information on temporal kinetics of BOLD signals ranging from the onset to the undershoot. We also add the sign of the time course without extracting any of the individual characteristics. Furthermore, we were able to use these ordinal scaled clustering results to jointly observe layer-wise differences in the signal processing across different studies. This is remarkable because due to differences in study protocols, which include different rat strains, anesthesia regimens, field strengths, temporal and spatial resolutions, this would hardly be possible with conventional methods. This highlights a further advantage of our analysis framework for stimulation-induced BOLD fMRI data, parallel to the recently reported protocol for functional connectivity during resting-state (Grandjean et al., 2023).

Yet, analyses of this type also come with challenges, such as the fact that clustering is not only dependent on the choice of metric, as we have explored. Decisions regarding the type of procedure or other parameters like the number of clusters must be preset. In addition, unsupervised machine learning is quickly unstable to data change (especially those with low signal responses). At least for parts of our analysis, we were able to prove the robustness of our preset version of HAC and its dependence on the SNR. Despite the robustness of this workflow, its transferability to other multi-dimensional datasets remains to be validated in future studies.

## Conclusion

In summary, we have developed a method to analyze task fMRI data in a completely data-driven manner. In this process, brain structures are determined by means of similarity relations of the BOLD response. A new Cluster Probability Profile (CPP) plot was introduced by spatially visualizing these structures over multiple measurements, illustrating the probability of a given signal character per voxel at the level of individual datasets or animals. We validated previously published study results using a different statistical analysis, which is entirely based on cluster distribution. Importantly, each cluster represents a group of parameters of the BOLD response instead of single parameters, which makes this procedure more robust. It is sensitive to detect differences in temporal BOLD kinetics of cortical modality processing and finally, we can effortlessly combine and compare datasets from different studies and imaging centers to extend statistical testing to larger cohort sizes. In addition, the described analysis pipeline can be used in the future for complex, time-dependent quantitative data sets, regardless of their data structure. This holds great potential for the future, as data-driven approaches will be increasingly used in future analysis routines.

## Supplemental Material

**Supplement Figure 1).**
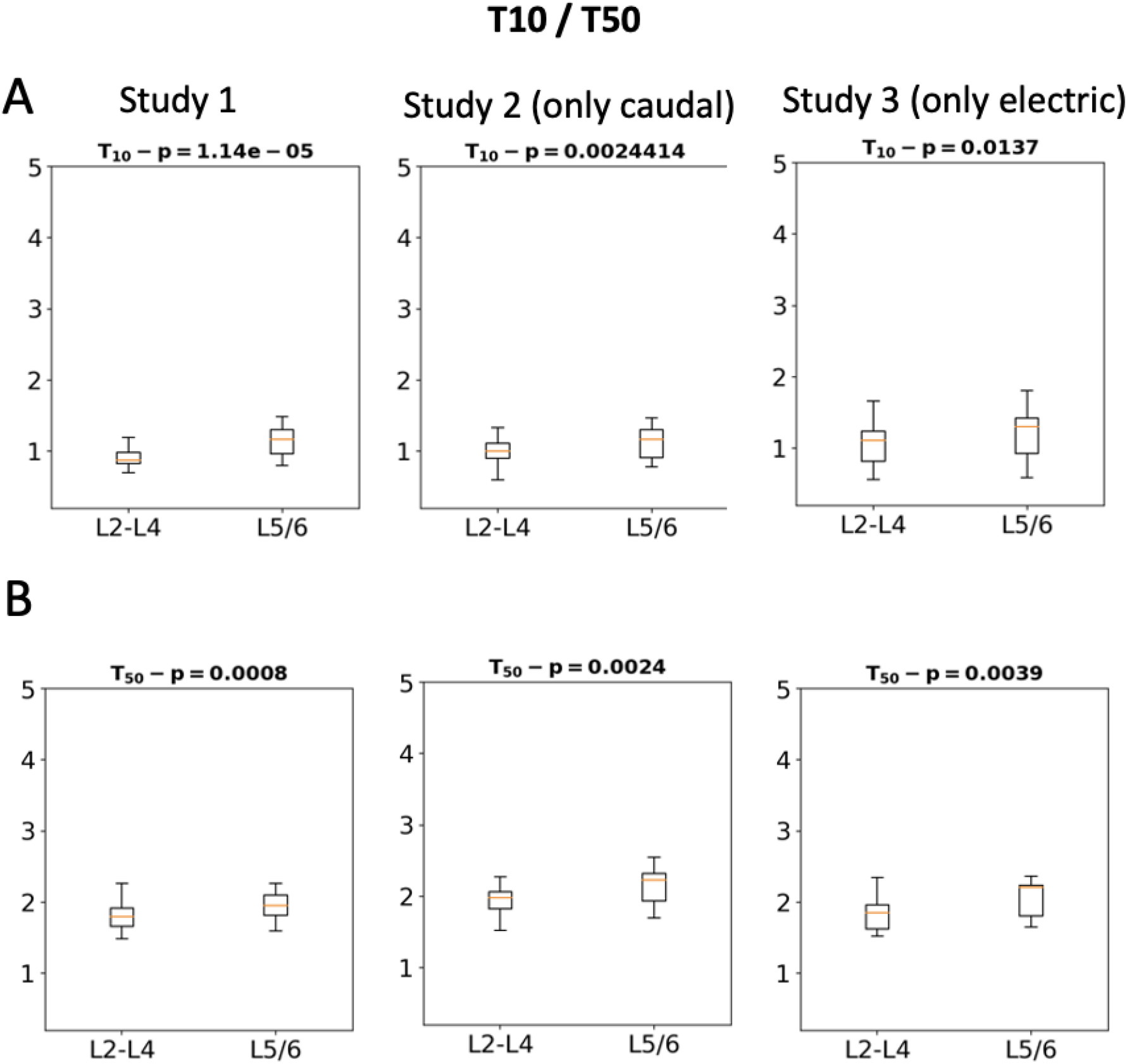
Comparison of T_10_ and T_50_ between LII/II+LIV and LV+LVI across all three studies.

